# Petite Integration Factor 1 (PIF1) helicase deficiency increases weight gain in Western diet-fed female mice without increased inflammatory markers or decreased glucose clearance

**DOI:** 10.1101/393355

**Authors:** Frances R. Belmonte, Nikolaos Dedousis, Ian Sipula, Nikita A. Desai, Aatur D. Singhi, Yanxia Chu, Yingze Zhang, Sylvie Bannwarth, Véronique Paquis-Flucklinger, Lea Harrington, Michael J. Jurczak, Robert M. O’Doherty, Brett A. Kaufman

## Abstract

Petite Integration Factor 1 (PIF1) is a multifunctional helicase present in nuclei and mitochondria. PIF1 knock out (KO) mice exhibit accelerated weight gain and decreased wheel running on a normal chow diet. In the current study, we investigated whether *Pif1* removal alters whole body metabolism in response to weight gain. PIF1 KO and wild type (WT) C57BL/6J mice were fed a Western diet (WD) rich in fat and carbohydrates before evaluation of their metabolic phenotype. Compared with weight gain-resistant WT female mice, WD-fed PIF1 KO females, but not males, showed accelerated adipose deposition, decreased locomotor activity, and reduced whole-body energy expenditure without increased dietary intake. Surprisingly, PIF1 KO females were protected against obesity-induced alterations in fasting blood glucose and glucose clearance. WD-fed PIF1 KO females developed mild hepatic steatosis and associated changes in liver gene expression that were absent in weight-matched, WD-fed female controls, linking hepatic steatosis to *Pif1* ablation rather than increased body weight. WD-fed PIF1 KO females also showed decreased gene expression of inflammatory markers in adipose tissue. Collectively, these data separated weight gain from inflammation and impaired glucose homeostasis. They also support a role for *Pif1* in weight gain resistance and liver metabolic dysregulation during nutrient stress.

## INTRODUCTION

The increasing prevalence of obesity is a global public health problem, which has been particularly striking among children and adolescents in both developed and developing countries [1]. Clinical complications of obesity include metabolic syndrome, which can progress to type 2 diabetes mellitus (T2DM) and cardiovascular disease. Current understanding of the genes involved in obesity is limited, despite its high heritability [2]. Approximately 90% of patients with T2DM are overweight or obese. However, while being overweight greatly increases the likelihood of developing T2DM [3], only a subset of obese individuals acquires T2DM. Factors such as tissue inflammation that drive obesity or that determine the diabetic phenotype in obese individuals are poorly understood.

Mouse models have been used to investigate genetic factors that alter the incidence and severity of obesity and diabetes in states of over nutrition, such as from a Western diet (WD) characterized by a high fat and carbohydrate content. We previously observed that male and female Petite Integration Frequency 1 helicase knockout (PIF1 KO) mice had increased body weight on regular chow [4]. This suggested that *Pif1* could play a role in the pathogenesis of obesity, although it remains unclear how *Pif1* contributes to body weight maintenance. PIF1 is a conserved ATP-dependent RNA/DNA helicase that localizes to both the nucleus and mitochondria [5]. In yeast, Pif1p inhibits telomerase [6-9], resolves short flaps in replication forks and unwinds secondary genomic structures that interfere with nuclear replication elongation [10, 11]. Mammalian PIF1 also interacts with telomerase. However, PIF1 KO mice exhibit normal telomere length after four generations of homozygosity [12], suggesting a different mechanism for this protein in metabolism.

The physiological basis for *Pif1* regulation of body weight has not been addressed. The primary objective of this study was to determine whether WD-fed PIF1 KO mice develop metabolic alterations under diet-induced weight gain. Toward that end, we measured major determinants of energy balance including locomotor activity, feeding, and energy expenditure by indirect calorimetry. Body composition, glucose tolerance and inflammation were also assessed. We found that PIF1 KO females, but not males, gained more weight on a WD, which was attributed specifically to increased fat mass. Surprisingly, PIF1 KO female mice were protected against the glucose intolerance and inflammation that are typically associated with obesity. The physiological basis for PIF1 KO female fat deposition was likely due to changes in physical activity. Our findings highlight an important role for a helicase such as PIF1 in regulating metabolism.

## MATERIALS AND METHODS

### Animals

Wild type C57BL/6J mice were from the Jackson Laboratory (Bar Harbor, ME). PIF1 KO mice were originally developed in the C57BL/6 background [4, 12], but the *Pif1* deletion was maintained in the C57BL/6J background exclusively for more than 10 generations. Animals were fed normal or WD chow *ad libitum* and housed in a 12 h light to dark cycling room. This study was performed in strict accordance with the recommendations in the “Guide for the Care and Use of Laboratory Animals” (2018) of the National Institutes of Health (Bethesda, MD) and was approved by the Institutional Animal Care and Use Committee of the University of Pittsburgh (Protocol Numbers: 15035458 and 18032212).

The experimental cohort for long-term WD studies was comprised of eight-to nine-week-old WT and PIF1 KO males and females exclusively fed a WD (41% fat, 27% sucrose, 19% protein, per calories; no. 96001; Harlan Teklad, Madison, WI) for 16 weeks. Four mice of the same sex, genotype and age were group housed together. Total body weight for all mice was measured weekly for 16 weeks. A separate experimental cohort for short-term WD studies consisted of nine-week-old, age- and weight-matched WT and PIF1 KO female mice (eight per group) that were singly housed to allow measurement of individual food intake and weight.

For the short-term WD feeding study, baseline metabolic measurements (as described in the following section) were obtained from female mice fed regular chow before transitioning to the WD. Metabolic measurements were repeated 14 days after WD initiation. Total body weight and WD intake were measured daily, Monday through Friday. Food intake was calculated for individual mice from the amount of food consumed per day in grams divided by the total body weight.

### Metabolism and body composition

Metabolic profiles of WT and PIF1 KO mice in both long- and short-term studies were compared using the Comprehensive Laboratory Animal Monitoring System (CLAMS, Columbus Instruments, Columbus, OH). Mice were acclimated for 24 h then energy expenditure, foraging activity, oxygen consumption and carbon dioxide production were measured for a second 24 h period. All animals underwent body composition analyses using ^1^H magnetic resonance spectroscopy (EchoMRI-100H, Houston, TX).

### Serum cytokine, diabetes, obesity, and cholesterol concentrations

WT and PIF1 KO females were fasted for 6 h prior to euthanasia. Blood was collected by cardiac puncture and centrifuged at 1000 x g for 10 min. Serum concentrations of cytokines (IL-1α, IL-1β, IL-12, eotaxin, G-CSF, IFN-γ, MCP-1, MIP-1β, TNF-α, IL-12p70), diabetes and obesity markers (ghrelin, gastric inhibitory polypeptide, glucagon-like peptide-1, glucagon, insulin, leptin, plasminogen activator inhibitor-1, resistin) were quantified with the Bio-Plex Pro Mouse Cytokine 23-plex and Diabetes 8-plex immunoassays, respectively (Bio-Rad, Hercules, CA). Total cholesterol, high-density lipoprotein and low-density lipoprotein/ very low-density lipoprotein serum concentrations were measured using colorimetric methods described in the manufacturer’s protocol (BioAssay Systems, Hayward, CA).

### Intraperitoneal glucose tolerance test

Six- and nine-month-old, regular chow-fed female WT and PIF1 KO mice and a separate cohort of 3.5- and five-month-old female WT and PIF1 KO mice were fasted for 6 h before measuring baseline glucose concentrations in whole blood sampled from a tail vein. D-glucose diluted in saline was administered at 1 g/kg body weight by intraperitoneal injection and blood glucose concentrations measured every 15 min for 2 h using Contour blood glucose test strips (Ascensia Diabetes Care US Inc, Parsippany, NJ) inserted into a glucometer.

### Liver pathology

Livers from female WT and PIF1 KO mice were fixed in 10% formalin for 24 h, transferred in ethanol and paraffin embedded before sectioning and staining. Liver sectioning and haematoxylin and eosin staining were performed at the Transplant Surgery Experimental Pathology Laboratory, Research Histology Services Core at the University of Pittsburgh. Liver histopathology was scored by a surgical pathologist blinded to the experimental cohort.

### Quantitative reverse transcription PCR

Total RNA was isolated [13] from powdered epididymal white adipose and liver tissues using TRIzol extraction (Invitrogen, Carlsbad, CA), and purified on a RNeasy column (Qiagen, Germantown, MD). Potential DNA contamination was removed by a DNA-free assay (ThermoFisher, Carlsbad, CA). Random hexamer primed reverse transcription was performed using the High Capacity cDNA synthesis kit as per the manufacturer’s protocol (ThermoFisher Scientific, Carlsbad, PA). Quantitative real time PCR was performed using PowerUp SYBR Green (ThermoFisher Scientific, Carlsbad, CA) on a QuantStudio 5 Real-Time PCR System (ThermoFisher Scientific, Carlsbad, CA). Reaction setup was performed using a Tecan Evo150 automated platform (Tecan, Morrisville, NC). Primers used in this study are presented in Table S1.

### Statistical analysis

Statistical analyses were performed using GraphPad Prism 7 software (GraphPad, La Jolla, CA). Student’s two-tailed, unpaired t-test was used to determine the significance of differences between two groups. Two-way repeated measures ANOVA with post-hoc Sidak’s multiple comparisons test were used to compare means obtained from metabolic cage data. Area under the concentration-time curve (AUC) analyses were performed to determine the total areas under the glucose tolerance test curves. P-values of <0.05 denoted significant differences. Standard error of the mean is shown for all experiments.

## RESULTS

### Female PIF1 KO mice gained weight and increased fat mass faster than WT mice when fed a WD

To further define the role of *Pif1* in regulating body weight, we fed male and female WT and PIF1 KO mice a WD for 16 weeks (characterized as long-term WD feeding) to induce weight gain. PIF1 KO and WT males gained weight rapidly at similar rates (Fig 1A), while body composition was similar in the two groups (Fig 1C). In contrast, PIF1 KO females gained significantly more weight and at a faster rate than WT females (Fig 1B). Body composition analyses demonstrated that these differences were due exclusively to increases in fat mass in WD-fed PIF1 KO females, with lean mass being the same in the two groups (Fig 1C). As the weight of WD-fed PIF1 KO females diverged from WT females, we focused exclusively on the female cohort in subsequent experiments.

**Fig 1.**
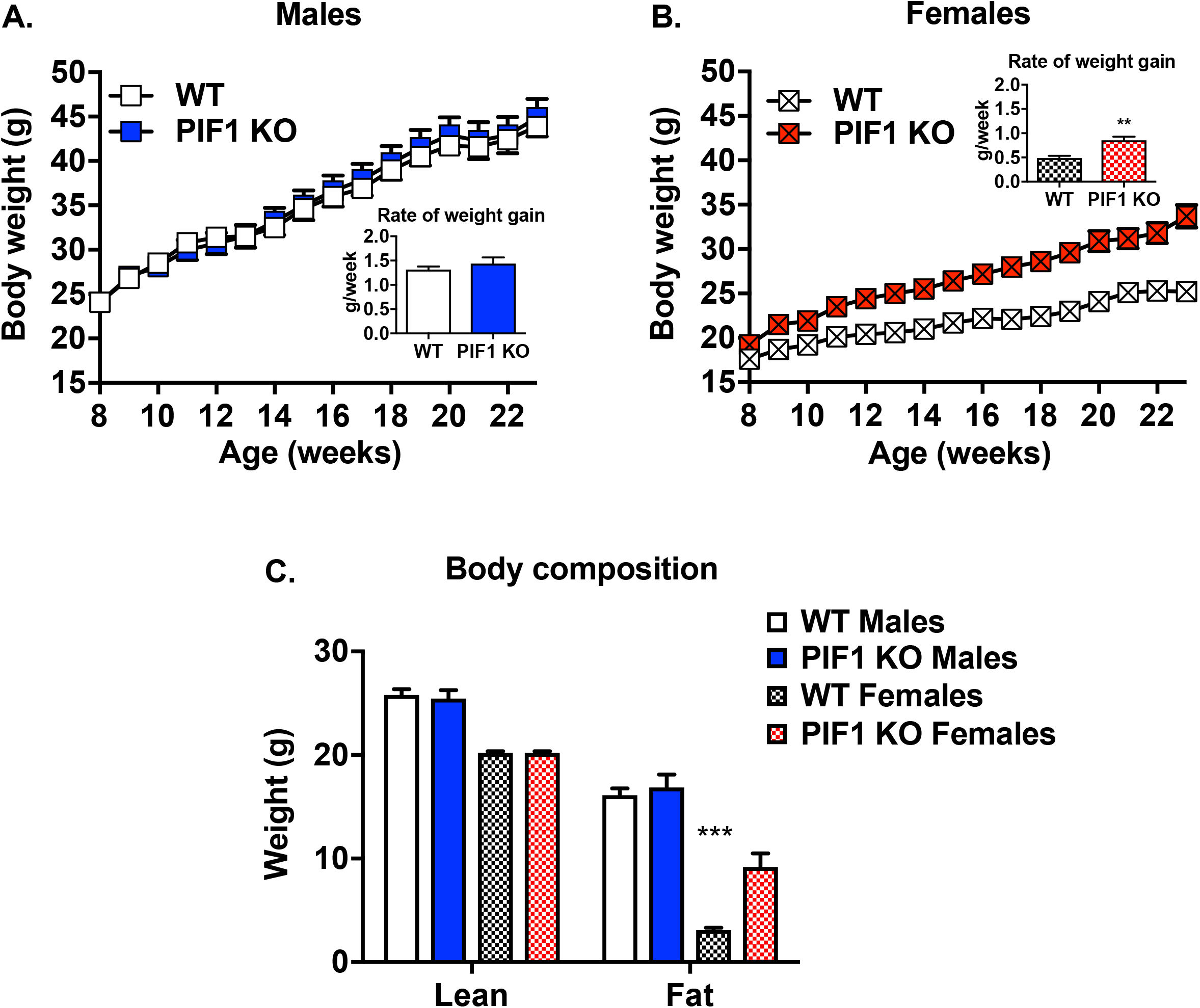
PIF1 knockout (KO) females, but not males, showed accelerated fat deposition compared with wild type (WT) mice during Western diet (WD) feeding. (A) Changes in body weight during 16 weeks of WD feeding in WT and PIF1 KO male, and (B) WT and PIF1 KO female mice. (C) Body composition as assessed through lean and fat mass measured in WT and PIF1 KO, male and female, 5-month-old mice after 12 weeks of WD feeding. Mean ± SEM; n = 8-12 mice per group; **p<0.01; ***p<0.001; g = grams.

### PIF1 KO female mice fed a WD for 16 weeks (long-term) showed decreased locomotor activity compared with WT female mice

To understand the physiological basis of body weight differences between WD-fed PIF1 KO and WT female mice, we measured the major determinants of energy balance. After four weeks of WD feeding, three-month-old PIF1 KO females showed reduced locomotor activity during both light and dark cycles and decreased whole-body energy expenditure across 24 h (Fig 2A, B). Following 12 weeks of WD feeding, 5-month-old PIF1 KO females continued to display decreased locomotor activity (Fig 2C), although WT animal activity also began to decrease with weight gain. Whole-body energy expenditure remained lower in 5-month-old PIF1 KO females during the dark cycle (Fig 2D). Substrate utilization, determined by the respiratory exchange ratio, was not different at any time point (data not shown).

**Fig 2.**
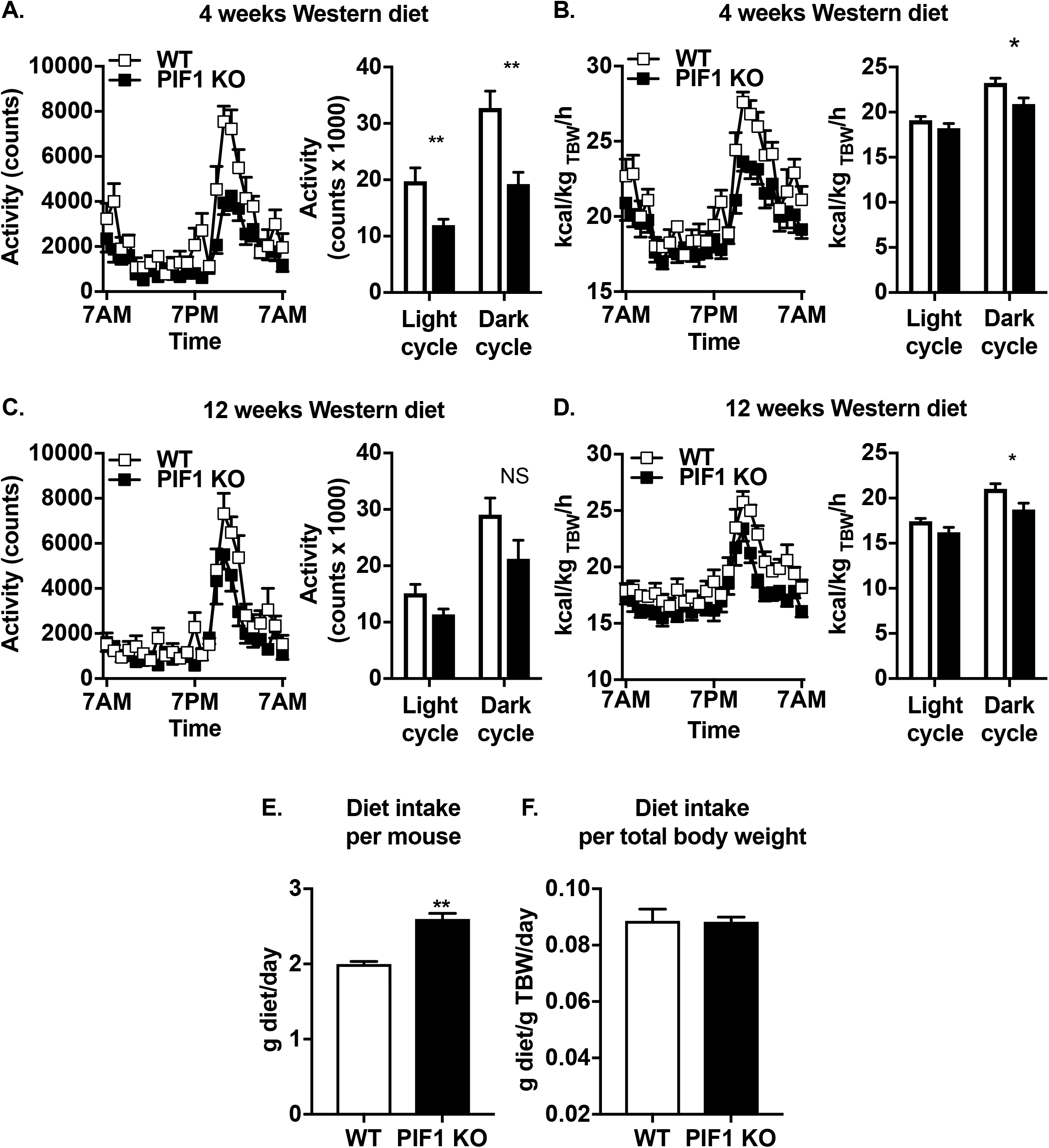
Long-term, Western diet (WD)-fed PIF1 KO mice exhibited decreased locomotor activity and whole-body energy expenditure. (A) Activity measured via beam breaks in 3-month-old, WD-fed PIF1 KO mice after 4 weeks of feeding, and (B) whole-body energy expenditure measured in 3-month-old, WD-fed PIF1 KO mice. (C) Locomotor activity measured in 5-month-old, WD-fed PIF1 KO mice after 12 weeks of feeding, and (D) whole-body energy expenditure measured in 5-month-old, WD-fed PIF1 KO mice. (E) WD food intake per mouse, (F) normalized to lean mass, and (G) normalized to total body weight in PIF1 KO mice after 16 weeks of feeding. Mean ± SEM; n = 8 mice per group; *p<0.05, **p<0.01; kcal/kg TBW/h = kilocalories per kilogram total body weight per hour; g = grams.

To assess whether decreased locomotion preceded weight gain, a separate cohort of age- and weight-matched WT and PIF1 KO female mice were fed a regular chow diet and then placed on a 14-day (short-term) WD diet.

Activity was measured in both dietary conditions. On regular chow, locomotor activity and whole-body energy expenditure were not different between groups (Fig S1A,B). After 14 days of WD, locomotor activity trended downward during the dark cycle in PIF1 KO females compared with WT females (Fig S1C), while energy expenditure did not differ between groups (Fig S1D). Thus, the decreased locomotor activity observed in the long-term WD-fed PIF1 KO females preceded the weight gain.

To determine whether WD was a potential driver for the weight gain phenotype in WD-fed PIF1 KO females, we assessed food consumption in the long- and short-term WD studies. Total WD intake per mouse (Fig 2E) and normalized to lean mass (Fig 2F) were higher overall in WD-fed PIF1 KO versus WT females. However, WD intake normalized to total body mass (Fig 2G) was similar between groups. Differences in body weight can confound interpretation of feeding data [14]. Therefore, we measured WD intake under body-weight matched conditions using short-term WD-fed animals and found no differences in feeding (Fig S1E). Thus, increased body weight in WD-fed PIF1 KO female mice was unlikely to be due to differences in food intake.

### Glucose tolerance is unchanged in WD-fed PIF1 KO females despite increased adiposity

As body weight strongly influences fasting blood glucose concentrations and glucose tolerance, we performed intraperitoneal glucose tolerance tests on regular chow- and WD-fed WT and PIF1 KO female mice.

Body weight-matched six-month-old (Fig 3A) and nine-month-old (Fig 3B) regular chow-fed females exhibited normal fasting blood glucose concentrations and glucose tolerance. Interestingly, despite the increased body weight and adiposity in long-term WD-fed PIF1 KO female mice (Fig 1), fasting blood glucose and glucose tolerance were not impaired relative to WT females after 6 and 12 weeks of WD-feeding (Fig 3C, D).

**Fig 3.**
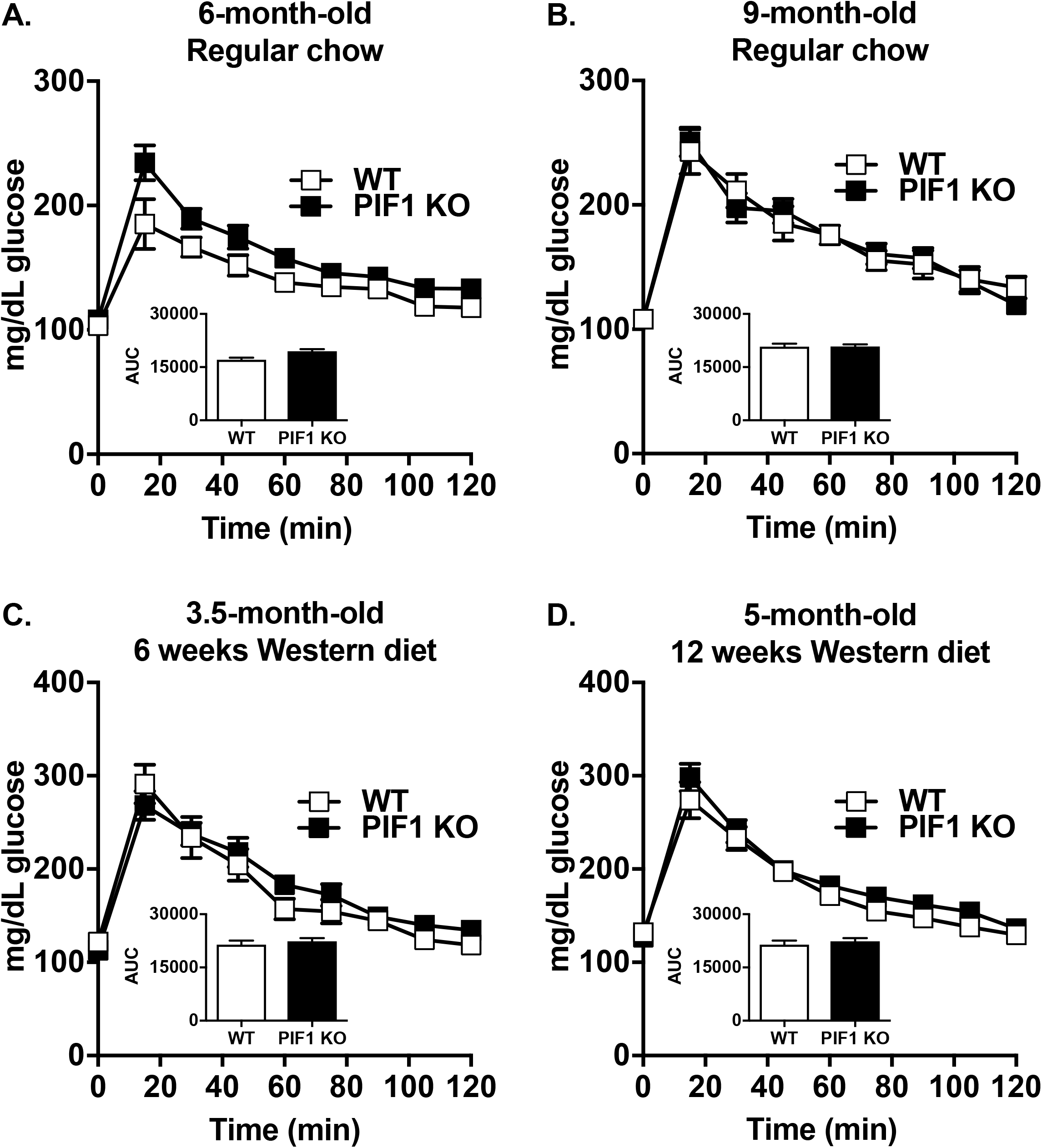
Western diet (WD) feeding did not affect blood glucose clearance in regular chow- and WD-fed PIF1 KO mice. (A) Glucose tolerance tests were performed on regular chow-fed, six-month-old WT and PIF1 KO mice, and on (B) the same mice at nine months of age. (C) Glucose tolerance tests conducted on 3.5-month-old WT and PIF1 KO mice after 6 weeks of WD feeding, and (D) 5-month-old WT and PIF1 KO mice after 12 weeks of WD feeding. Mean ± SEM; n = 8-12 mice per group; mg/dl = milligrams per deciliter; AUC = area under the curve.

The apparent protection against insulin resistance prompted us to examine specific serum factors that are commonly dysregulated during diabetes and obesity (Table 1). Consistent with normal glucose tolerance, fasting insulin concentrations were similar between genotypes after long-term WD feeding. As expected with the increased adiposity in the PIF1 KO mice, leptin concentrations were significantly elevated in serum. Other factors associated with diabetes and obesity including ghrelin, gastric inhibitory polypeptide (GIP), glucagon-like peptide-1 (GLP-1), plasminogen activator inhibitor-1 (PAI-1), resistin, cholesterol, high-density lipoprotein (HDL) and low-density lipoprotein/ very low-density lipoprotein (LDL/VLDL) were largely unchanged. Glucagon, a metabolic hormone that is secreted during fasting and is a positive regulator of gluconeogenesis, was upregulated. This likely occurs to maintain normoglycemia in obese animals in the fasted state [15].

**Table 1.**
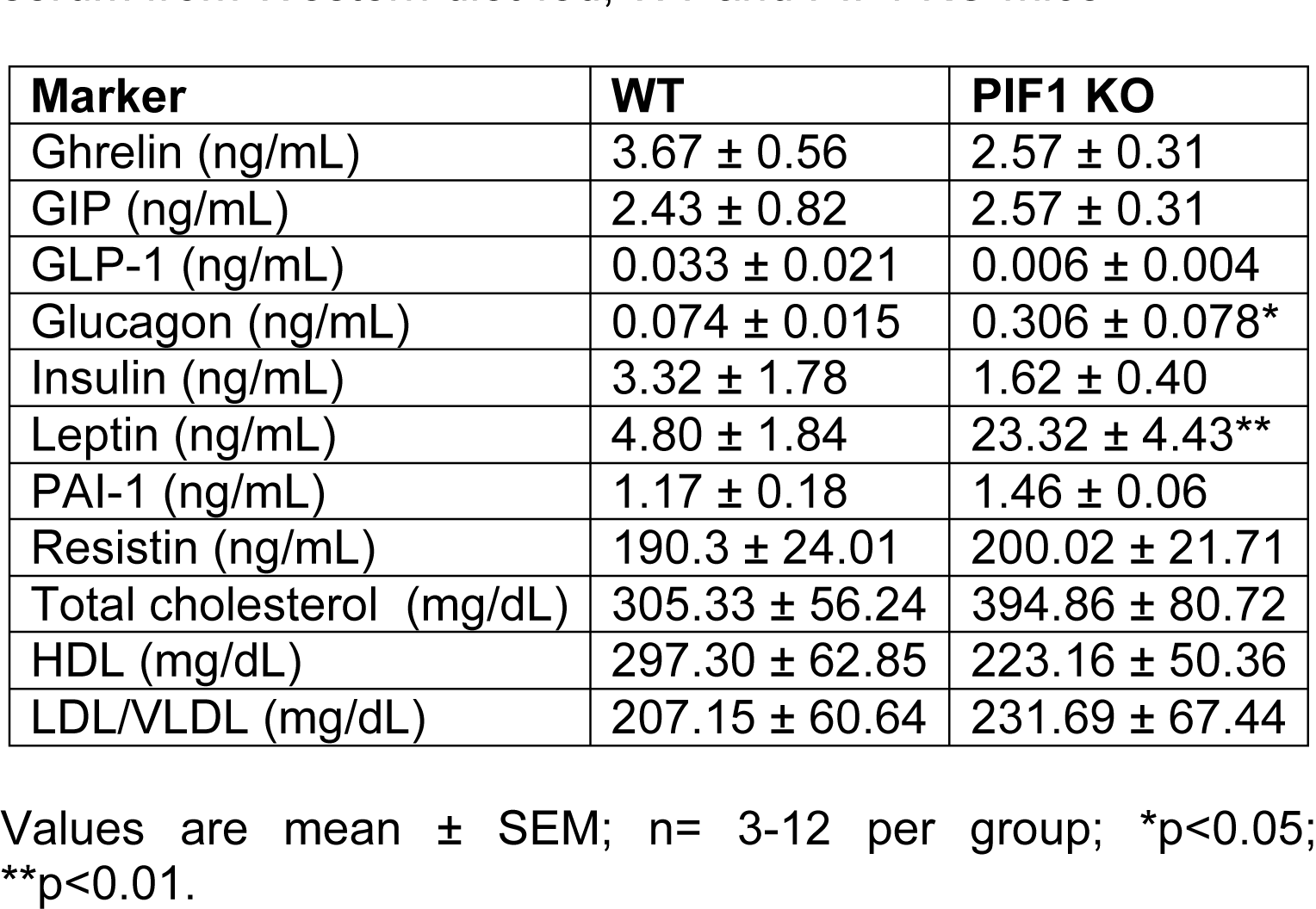
Diabetes, obesity, and cholesterol concentrations in serum from Western diet-fed, WT and PIF1 KO mice

### WD-fed PIF1 KO female mice exhibited mild hepatic steatosis with modest alterations in markers of gluconeogenesis and mitochondrial biogenesis

During weight gain, hepatic fat accumulation can lead to alterations in fatty acid metabolism and cholesterol handling. To investigate whether PIF1 KO mice were particularly vulnerable to hepatic steatosis, haematoxylin and eosin stained liver sections from the long-term WD cohort were evaluated in a blinded fashion by a liver pathologist. PIF1 KO mice uniformly exhibited mild hepatic steatosis compared to WT mice (Fig 4A). To determine whether differences in steatosis were due to obesity, we compared a weight-matched subset of animals and found steatosis only in the PIF1 KO animals. These data indicate that the hepatic steatosis observed in WD-fed PIF1 KO mice is independent of weight gain.

**Fig 4.**
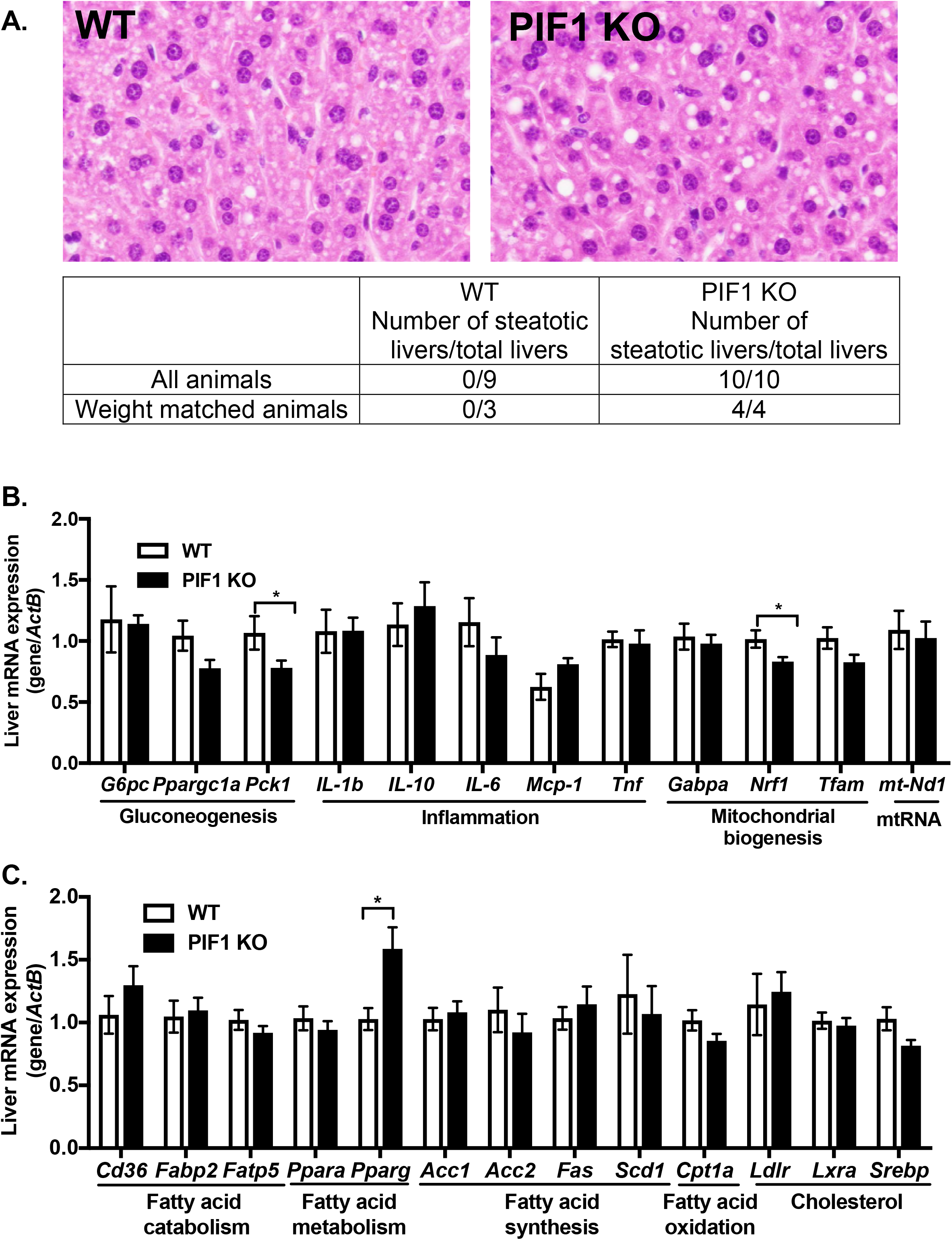
Livers from long-term, Western diet (WD)-fed PIF1 KO mice showed minor macro hepatic steatosis, and transcripts for steatosis and gluconeogenesis markers were altered in the PIF1 KO livers. (A) Representative images of hematoxylin and eosin stained livers from WD-fed WT and PIF1 KO females (n=6 mice per group). The table below Fig 4A shows the total number of livers from all animals and weight-matched, WT and PIF1 KO livers that were blindly scored for the presence of macro hepatic steatosis. (B) Liver cDNA from WD-fed, WT and PIF1 KO mice was used to measure the expression of genes involved in gluconeogenesis, inflammation, mitochondrial biogenesis, mitochondrial RNA, and (C) fatty acid catabolism, fatty acid metabolism, fatty acid synthesis, fatty acid oxidation and cholesterol. mRNA expression relative to beta actin (*ActB*) reference gene in liver tissue. Mean ± SEM; n = 8-12 mice per group; *p<0.05.

To examine differences in the transcriptional response between long-term WD-fed WT and PIF1 KO livers, we analyzed the expression of a set of genes with established roles (gluconeogenesis, inflammation, mitochondrial biogenesis and lipid metabolism) in liver fat accumulation (Fig 4B, C). Transcripts for phosphoenolpyruvate carboxylkinase 1 (*Pck1*), a rate-limiting enzyme for gluconeogenesis, and nuclear respiratory factor 1 (*Nrf1*), a transcription factor involved in regulating mitochondrial biogenesis, were decreased in livers from WD-fed PIF1 KO mice (Fig 4B). Expression of additional genes known to regulate gluconeogenesis (*G6pc, Ppargc1a*), inflammation (*Il-1b, Il-10, Il-6, Mcp-1, Tnf*), mitochondrial biogenesis (*Gabpa, Tfam)* or mitochondrial DNA encoded genes (*mt-Nd1*) were similar between groups (Fig 4B). Additionally, we did not observe any differences in mitochondrial DNA (mtDNA) copy number or mitochondrial complex I activity between WD-fed WT and PIF1 KO female livers (data not shown). PIF1 KO livers showed increased expression of peroxisome proliferator-activated receptor gamma (*Pparg*) (Fig 4C), consistent with the steatotic phenotype [16-21]. Other genes that regulate fatty acid and cholesterol metabolism (*Cd36, Fabp2, Fatp5, Ppara, Acc1, Acc2, Fas, Scd1, Cpt1a, Ldlr, Lxra, and Srebp*) were unchanged in WT and PIF1 KO livers. Overall, these results showed that WD-fed PIF1 KO female mice developed mild hepatic steatosis with modest gene expression changes in relevant metabolic pathways.

### Adipose tissue from WD-fed PIF1 KO females has decreased mRNA expression of inflammatory markers without changes in markers of adiposity, fatty acid metabolism and mitochondrial biogenesis

As fat mass was increased in WD-fed PIF1 KO females compared with WD-fed WT mice, we assessed whether key markers of obesity pathogenesis were altered in epididymal white adipose tissue. No differences in mRNA expression of selected adiposity, fatty acid metabolism, mitochondrial biogenesis or mitochondrial RNA genes (*Adipoq, Dlk1, Lep, Pparg, Gabpa, Nrf1, Tfam*, and *mt-Nd1*) were observed between groups (Fig 5). Interestingly, despite the increased adiposity in WD-fed PIF1 KO females, which is commonly associated with adipose tissue inflammation [22-26], mRNA expression of *Mcp-1* was four-fold downregulated in PIF1 KO mice, whereas other inflammatory genes (*Il-10, Il-1b, and Tnf*) trended towards lower expression in adipose tissue (Fig 5). To examine systemic inflammation, we measured serum concentrations of pro-inflammatory cytokines, but detected no differences between PIF1 KO and WT mice fed a long-term WD (Table 2).

**Table 2.** Cytokines measured in serum from Western diet-fed, WT and PIF1 KO mice

| **Cytokine** | **WT** | **PIF1 KO** |
| --- | --- | --- |
| IL-1α (ng/mL) | 9.9 ± 2.0 | 8.0 ± 1.6 |
| IL-1β (ng/mL) | 206.2 ± 60.6 | 124.2 ± 23.4 |
| IL-12 (ng/mL) | 423.9 ± 98.2 | 695.2 ± 85.7 |
| Eotaxin (ng/mL) | 545.6 ± 143.6 | 524.1 ± 82.0 |
| G-CSF (ng/mL) | 31.5 ± 23.4 | 42.6 ± 10.2 |
| IFN-γ (ng/mL) | 20.6 ± 6.1 | 21.2 ± 3.4 |
| MCP-1 (ng/mL) | 166.4 ± 41.2 | 134.1 ± 34.5 |
| MIP-1β (ng/mL) | 29.7 ± 1.9 | 21.5 ± 3.6 |
| TNF-α (mg/dL) | 65.5 ± 18.6 | 52.3 ± 12.8 |
| IL-12p70 (mg/dL) | 49.4 ± 16.4 | 42.9 ± 9.1 |

Values are mean ± SEM; n= 3-12 per group.

**Fig 5.**
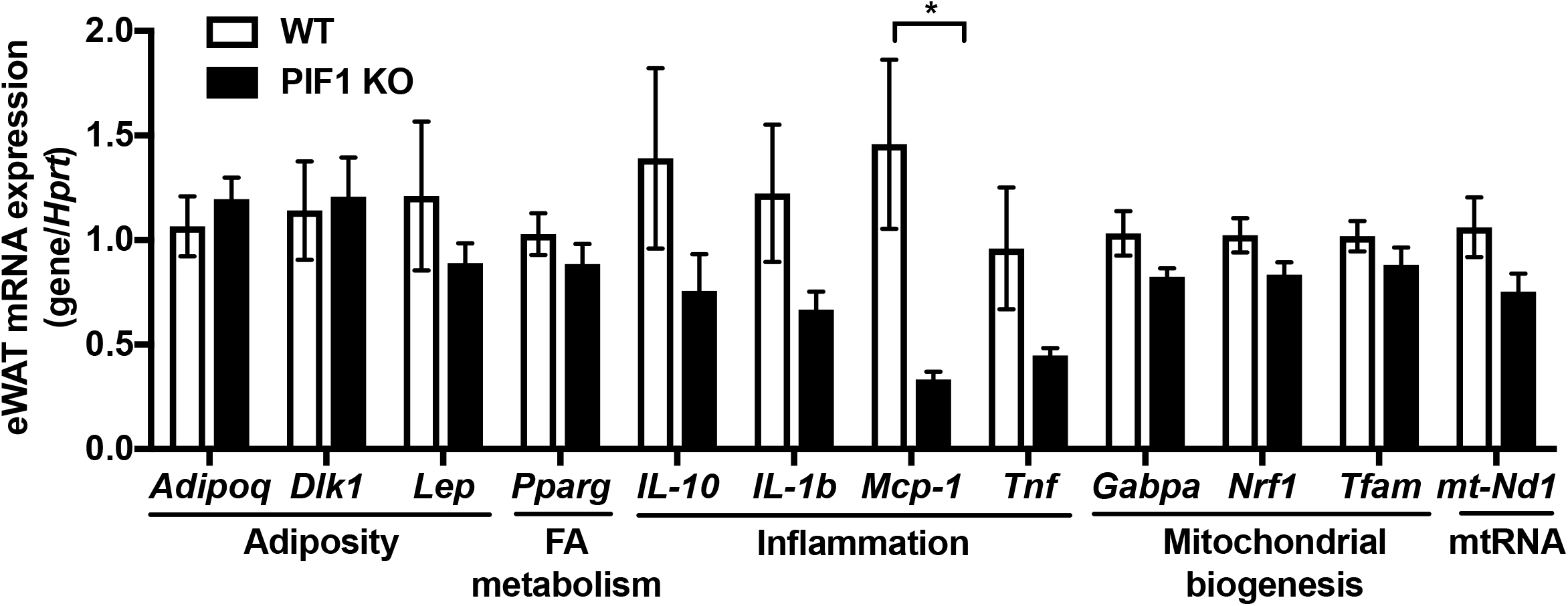
Transcripts for inflammatory markers were decreased in adipose tissue from Western diet (WD)-fed PIF1 KO females. Epididymal white adipose tissue (eWAT) cDNA from WD-fed, WT and PIF1 KO mice was used to measure transcripts for genes involved in adiposity, fatty acid (FA) metabolism, inflammation, mitochondrial biogenesis and mitochondrial RNA. mRNA expression relative to the hypoxanthine-guanine phosphoribosyl transferase (*Hprt*) reference gene. Mean ± SEM; n = 8-12 mice per group; *p<0.05.

## DISCUSSION

While we have previously shown that PIF1 KO mice gain weight on a regular chow diet, it was not known whether PIF1 KO mice develop metabolic alterations under WD-induced weight gain. WD-fed PIF1 KO females had increased body weight and fat mass, but, paradoxically, similar fasting blood glucose concentrations and glucose tolerance compared with WD-fed WT females. Interestingly, the PIF1 KO females also had decreased expression of adipose tissue inflammatory markers. WD-fed PIF1 KO females developed a mild hepatic steatosis that appeared independent of weight gain, and without severe gene expression derangement.

The finding that WD-fed PIF1 KO females diverged in weight from WT females, while the males gained weight at a similar rate, was unexpected. The C57BL/6J strain used to generate the PIF1 KO mice has a sex-specific response to WD feeding [27, 28]. Specifically, female C57BL/6J mice gain less weight and are generally better protected against WD-induced metabolic syndrome than males [29]. Thus, our findings suggest that WD-fed PIF1 KO females are susceptible to diet-induced weight gain or lose protection against weight gain. The dissociation between obesity and metabolic dysfunction in WD-fed PIF1 KO female mice is reminiscent of metabolically healthy obesity [30, 31]. The decreased expression adipose tissue inflammatory markers in WD-fed PIF1 KO females compared with WD-fed WT controls may account for some aspects of the healthy obese phenotype, but requires further investigation. Similarly, the absence of changes in fasting blood glucose concentrations and glucose tolerance despite increased hepatic steatosis in WD-fed PIF1 KO females was surprising given the strong, positive association between liver fat, hyperglycemia and hepatic insulin resistance [32-34]. Future studies using isotopic infusion to measure rates of hepatic glucose production and insulin sensitivity directly, such as the hyperinsulinemic-euglycemic clamp, will be necessary to resolve this apparent paradox.

In addition to PIF1 helicase, experiments have been conducted in transgenic *Twinkle* mtDNA replication helicase mice [35]. TWINKLE is a nuclear-encoded, mtDNA replication helicase where specific mutations can cause autosomal dominant progressive external ophthalmoplegia (adPEO), a neurological disease associated with mtDNA deletions causing progressive neuropathy with sensorineural loss [36]. Expression of a transgenic copy of an adPEO *Twinkle* allele in mice causes the accumulation of mtDNA deletions in the muscle and brain, now known as “Deletor” mice. Despite the presence of mtDNA deletions, Deletor mice do not display decreased physical performance or increased body weight [35]. These results from the Deletor mice suggest that the mtDNA deletions in the skeletal muscle of PIF1 KO mice previously described [4] do not contribute to the decreased activity and increased body weight shown in the current study.

The likely mechanism of increased weight gain in WD-fed PIF1 KO females was decreased locomotor activity. Both light and dark cycle activities were significantly reduced in PIF1 KO female mice after just four weeks of WD feeding when body weight differences were apparent, but modest. Additionally, there was a trend of decreased activity (25% reduction) after just two weeks of WD feeding prior to gross differences in body weight (Fig S1C). The importance of this two-week time point is that it was a weight-matched (pre-weight gain) cohort [37] that was evaluated pre- and post-WD exposure. Notably, when the weight trend is analyzed across all experiments and timepoints in the study, we find a significant difference between WT and PIF1 (Fig S1E, F). In the absence of pre-weight divergence hyperphagia, the evidence best aligns with the conclusion that there is an interaction between PIF1 deficiency and Western diet that modulates locomotor activity to accelerate weight gain. By extension, we speculate that a PIF1 deficiency in humans might only cause obesity on Western diets.

The current study improves upon our prior understanding of weight gain in PIF1 KO mice [4], showing a dietary influence and sex-based differential response, but there are multiple outstanding questions remaining. We do not know the tissues or cell types that are crucial for regulating this interaction between PIF1 deficiency and Western diet, which could be limited to the central nervous system, liver, skeletal muscle, or thermal regulation (i.e. brown fat). We do not understand the reason for the sex-based response and potential role of estrogen signaling in this process. Furthermore, we do not know whether the key function of PIF1 in the phenotypes reported here are due to the nuclear or mitochondrial functions of PIF1. These questions highlight the need for new mouse models, such as tissue specific PIF1 knock-outs and nuclear and mitochondrial PIF1 knock-outs to further interrogate the role of this helicase in metabolism.

## Acknowledgements

BAK was supported by NIGMS (R01GM110424). RMO was supported by NIDDK (R01DK102839) and the Center for Metabolism and Mitochondrial Medicine (C3M) funding. FB was supported by NIDDK (T32DK007052) and the animal costs were supported by NIGMS and Stimulating Pittsburgh Research in Geroscience (SPRIG) Award. ND was supported by C3M. ADS was supported by Pittsburgh Liver Research Center (PLRC) at the University of Pittsburgh. The content of this manuscript is solely the responsibility of the authors and does not necessarily represent the official views of the National Institutes of Health.

---

**Fig S1.**
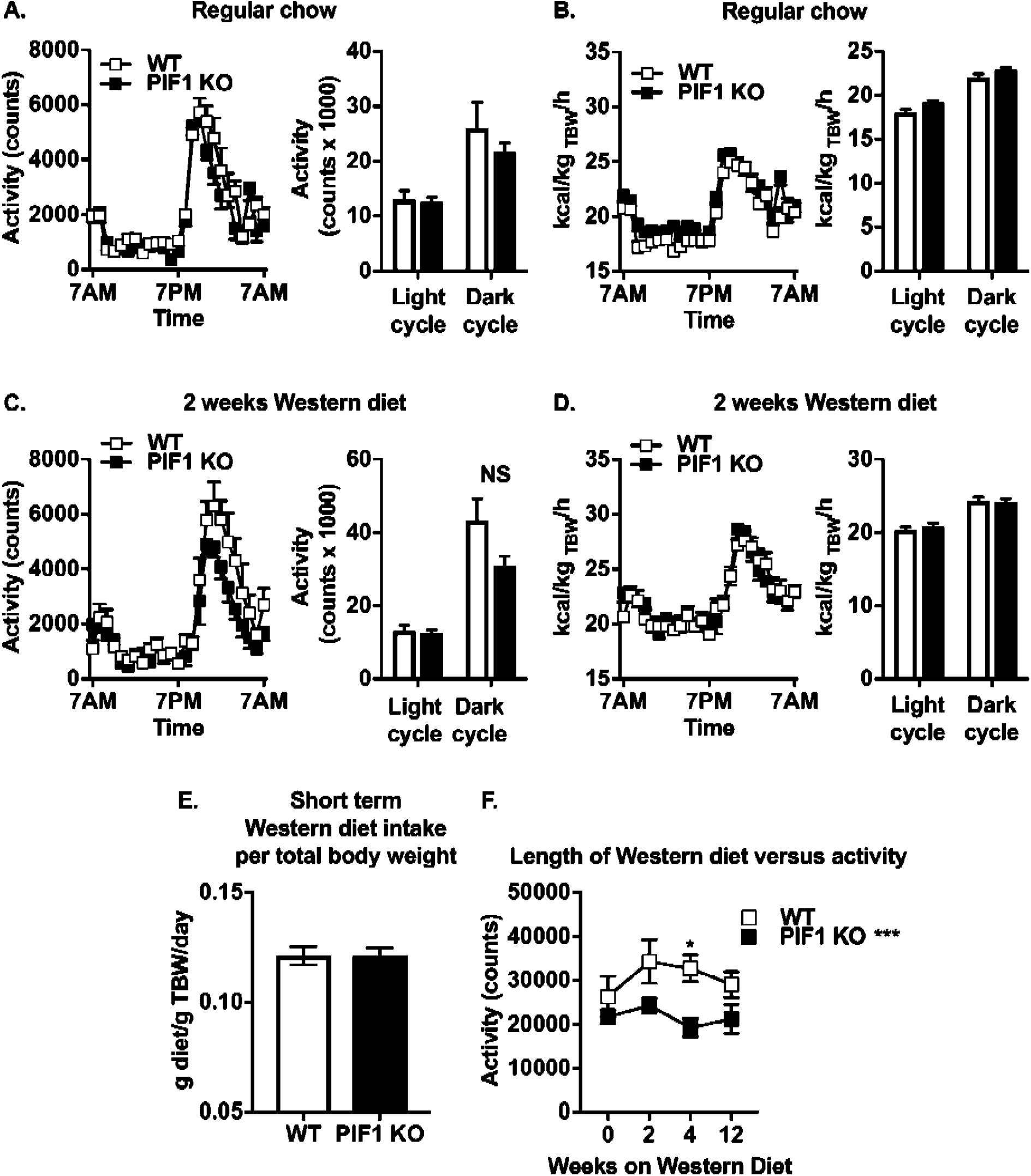
Locomotor activity from short-term. Western diet (WD)-fed PIF1 KO mice was decreased prior to weight divergence between WD-fed WT and WD-fed PIF1 KO females. (A) Locomotor activity, and (B) whole-body energy expenditure were measured in 2-month-old, regular chow-fed WT and PIF1 KO females. A separate cohort of WD-fed WT and PIF1 KO females were placed on a short-term WD for 2 weeks to measure (C) locomotor activity, (D) whole-body energy expenditure, and (E) food intake. (F) Locomotor activity of the short- and long-term WD cohorts over the course of 12 weeks. Mean ± SEM; n = 8 mice per group; *p<0.05; kcal/h = kilocalories per hour; kg TBW = kilograms total body weight.

**Table S1.**
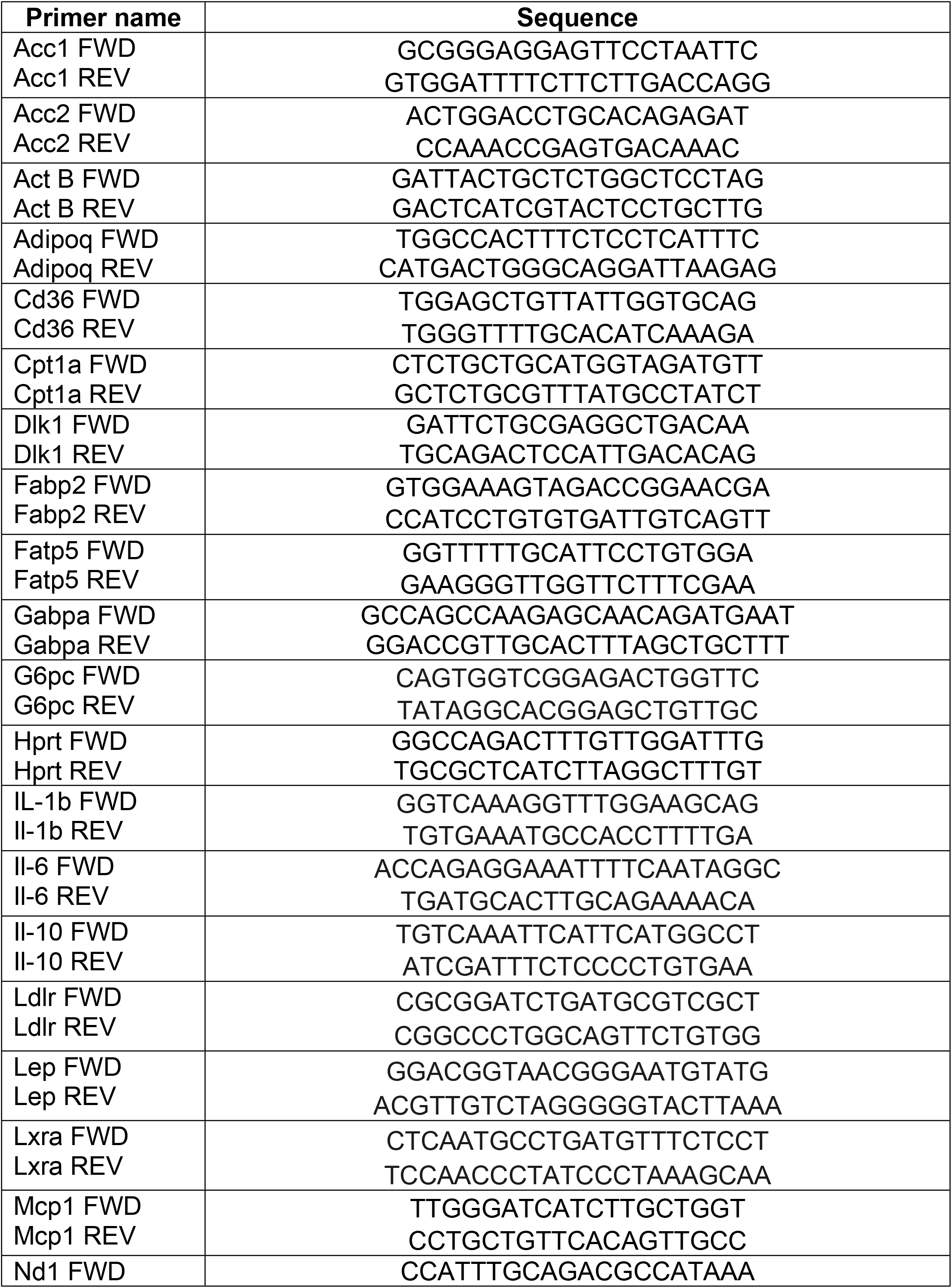
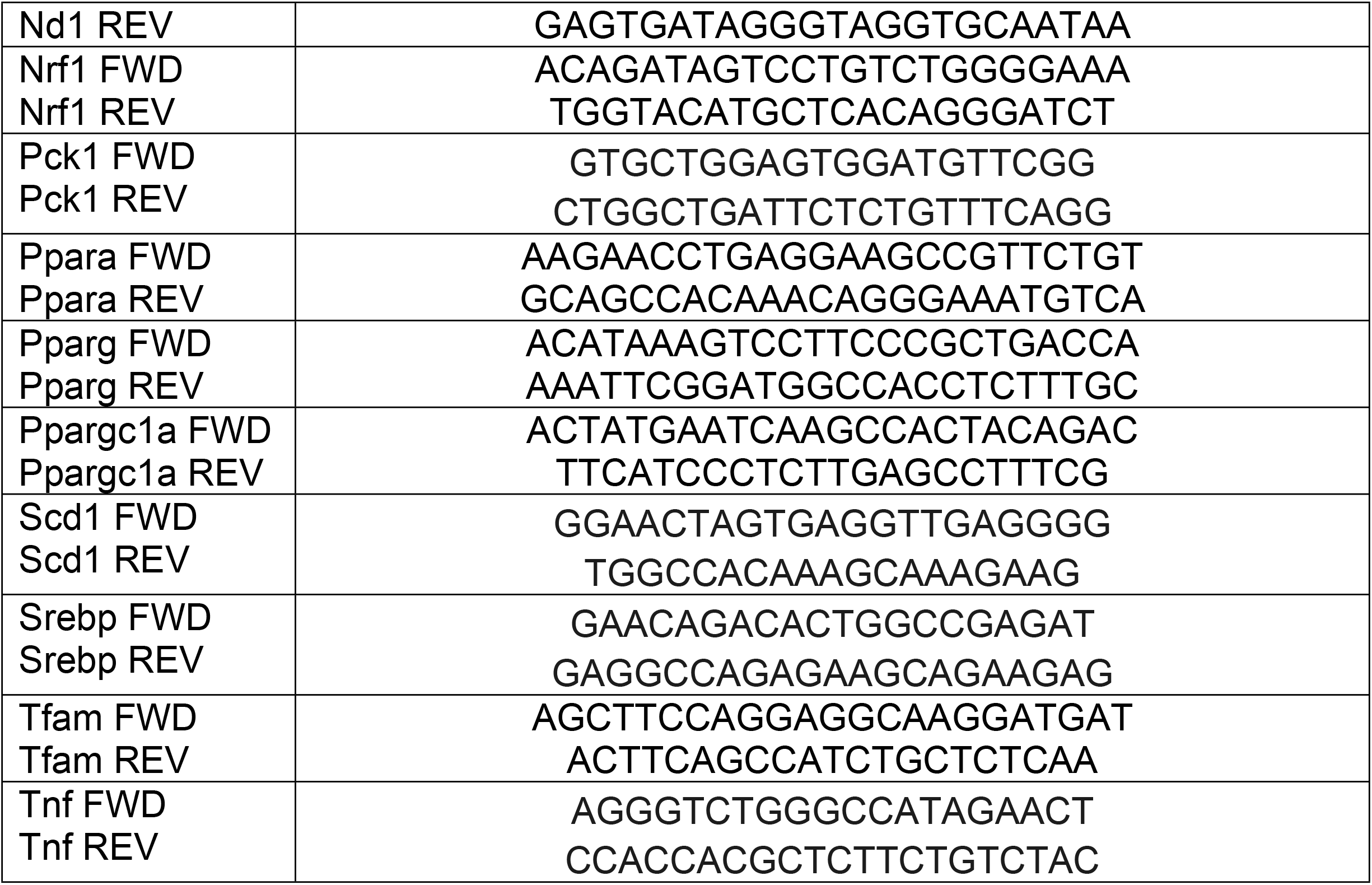
List of primers used to measure gene expression in RNA isolated from white adipose tissue and liver

